# Brain Connectivity Tracks Effects of Chemotherapy Separately from Behavioral Measures

**DOI:** 10.1101/352690

**Authors:** Omid Kardan, Mary K. Askren, Misook Jung, Scott Peltier, Bratislav Misic, Nathan W. Churchill, Patricia A. Reuter-Lorenz, Bernadine Cimprich, Marc G. Berman

## Abstract

Several studies in cancer research have suggested that cognitive dysfunction following chemotherapy, referred to in lay terms as “chemobrain”, is a serious problem. At present, the changes in integrative brain function that underlie such dysfunction remains poorly understood. Recent developments in neuroimaging suggest that patterns of functional connectivity can provide a broadly applicable neuromarker of cognitive performance and other psychometric measures. The current study used multivariate analysis methods to identify patterns of disruption in resting state functional connectivity of the brain due to chemotherapy and the degree to which the disruptions can be linked to behavioral measures of distress and cognitive performance. Sixty two women (22 healthy control, 18 patients treated with adjuvant chemotherapy, and 22 treated without chemotherapy) were evaluated with neurocognitive measures followed by self-report questionnaires and open eyes resting-state fMRI scanning at three time points: diagnosis (M0, pre-adjuvant treatment), at least 1 month (M1), and 7 months (M7) after treatment. The results indicated deficits in cognitive health of breast cancer patients immediately after chemotherapy that improved over time. This psychological trajectory was paralleled by a disruption and later recovery of resting-state functional connectivity, mostly in the parietal and frontal brain regions. The functional connectivity alteration pattern seems to be a separable treatment symptom from the decreased cognitive health. More targeted support for patients should be developed to ameliorate these multi-faceted side effects of chemotherapy treatment on neural functioning and cognitive health.

## Introduction

Adjuvant chemotherapy is a life-saving procedure in breast cancer treatment. However, considerable evidence suggests that cognitive dysfunction following chemotherapy, referred to in lay terms as “chemobrain,” is a serious mental health issue that is poorly understood (Bernstein, McCreath, Komeylian, & Rich, 2017; Jung et al., 2017; Schagen & Wefel, 2017; Wefel, Vardy, Ahles, & Schagen, 2011). Recent studies in network neuroscience have shown that relatively sparse patterns of functional connectivity strength during rest can signify broadly applicable neuromarkers of cognitive measures such as sustained attention (Rosenberg et al., 2016) and other psychometric and behavioral measures, thus linking resting state functional connectivity to cognition (e. g., (Biazoli et al., 2017; Smith et al., 2015). Given the established utility of resting-state neuroimaging in characterizing the neural basis of cognitive (dys)function, we used resting-state Blood-Oxygenation Level Dependent functional Magnetic Resonance Imaging (BOLD fMRI) to investigate changes in functional connectivity pre-to post-treatment and after recuperation, and its relationship with objective and subjective measures of cognitive functioning and health in women diagnosed with breast cancer.

There is ample evidence that chemotherapy is associated with cognitive deficits, and that some deficits are independent of cancer diagnosis and treatment and the accompanying distress (see (Schagen & Wefel, 2017) for a review). However, some studies have linked pre-treatment distress, worry, and fatigue associated with cancer diagnosis and treatment related worry to decreased functional connectivity of the BOLD signal in brain regions associated with attention and memory even before chemotherapy has begun (Andryszak, Wiłkość, Izdebski, & Żurawski, 2017; Churchill Nathan W. et al., 2015; Reuter-Lorenz & Cimprich, 2013). Nevertheless, researchers have yet to identify brain connectivity changes that are due to direct effects of chemotherapy versus connectivity changes that follow the cognitive and emotional distress associated with diagnosis and treatment (Ahles et al., 2010; Askren et al., 2014; Berman, Askren, et al., 2014; Cimprich et al., 2010; Dumas et al., 2013; Menning et al., 2015; Wefel Jeffrey S., Saleeba Angele K., Buzdar Aman U., & Meyers Christina A., 2010). Understanding the brain changes that explain the contribution of overall symptom burden (Jung et al., 2017) to ‘chemobrain’ is an essential step towards achieving effective interventions before, during, and after chemotherapy in order to more effectively restore cognitive functioning in cancer survivors.

In the current study, we used multivariate analysis methods to identify the pattern of disruption in resting state functional connectivity of the brain due to chemotherapy and the degree to which those patterns of disruption can be linked to measures of distress and cognitive performance. Here we quantify: 1) the effect of cancer treatment on resting-state functional connectivity, 2) the effect of cancer treatment on objective and subjective measures of cognitive performance and well-being, and 3) the relationship between resting-state functional connectivity and objective and subjective psychological measures. All of these tests are performed in a longitudinal manner, i.e., testing the effects at three timepoints: before, during and after treatment. These results will have important implications for how to treat the psychological and physiological effects on the brain that accompany breast cancer treatment and improve recovery from a psychological and physiological standpoint.

## Methods

### Participants

Sixty six right-handed women were recruited for 3 sessions from the University of Michigan Comprehensive Cancer Center, including two groups of women surgically treated for breast cancer (stage 0 - IIIa) awaiting adjuvant chemotherapy (CT, n = 22) or radiation-therapy without chemotherapy (non-CT, n = 22) and age-matched healthy controls (HC, n = 22) with negative mammograms that occurred within a year. Four women of the CT group did not return for post-baseline assessments due to unstable medical conditions or new MRI contraindications, resulting in n = 18 patients from the CT group to have data for all 3 time points. Screening criteria included absence of cognitive disorder (Mini-Mental Status Examination) (Folstein, Folstein, & McHugh, 1975), clinical depression (Patient Health Questionnaire, PHQ-8) (Kroenke et al., 2009), and secondary diagnosis of neurological or psychiatric disorders. The University of Michigan Institutional Review Board for Medicine approved all the studies, and all participants provided written informed consent.

### Design

Participants were evaluated with neurocognitive measures followed by self-report questionnaires and open eyes resting-state fMRI scanning at three time points. Assessments occurred at: diagnosis (M0, pre-adjuvant treatment), at least 1 month (M1), and 7 months (M7) after completion of treatment, resulting in 186 completed fMRI scanning sessions.

### Self-reported measures and objective tasks

Cognitive health of the participants was measured using objective cognitive performance and subjective emotional and cognitive well-being measures (i. e., behavioral variables). Subjective assessments included cognitive complaints (Attentional Function Index, AFI), physical symptom severity (Breast Cancer Prevention Trial symptom scales, BCPT) (Cella et al., 2008), psychological distress-worry (Three-Item Worry Index, TIWI) (Kelly, 2004), anxiety (State-Trait Anxiety Inventory, STAI) (Spielberger Charles D., 2010), depression (PHQ-8) (Kroenke et al., 2009), fatigue (Functional Assessment of Cancer Therapy: Fatigue, FACT-F) (Yellen, Cella, Webster, Blendowski, & Kaplan, 1997), and sleep problems (Pittsburgh Sleep Quality Index, PSQI) (Buysse, Reynolds, Monk, Berman, & Kupfer, 1989).

Objective cognitive assessments included the Verbal Working Memory Task (VWMT) that occurred during a separate fMRI scanning (Askren et al., 2014; Berman, Askren, et al., 2014; Nelson, Reuter-Lorenz, Sylvester, Jonides, & Smith, 2003), and two Digit Span tasks: Backwards (DSB) and Forward (DSF) (Gerton et al., 2004; Reynolds, 1997). For each trial (total of 192) in the VWMT, participants were presented with a set of four letters for 1,500 ms. Following a 3,000 ms delay interval, they were presented with a “probe” letter for 1,500 and asked whether it was a member of the current memory set. This task was done during four runs of fMRI scanning for which the brain data were reported in a previous study (Jung et al., 2017). For the current study we used the reaction time (RT) and accuracy measures of the WVMT.

### fMRI acquisition parameters

Images were acquired on a GE SIGNA 3 Tesla scanner, equipped with a standard quadrature head coil. Functional T2* weighted images were acquired using a spiral sequence with 25 contiguous slices with 3.75 × 3.75 × 5 mm voxels with repetition time (TR) = 1500 ms; echo time (TE) = 30 ms; flip angle =70°; field of view (FOV) = 24 cm for 360 seconds of rest (eyes open). A T1-weighted gradient echo anatomical overlay was acquired using the same FOV and slices (TR = 225 ms, TE = 5.7 ms, flip angle = 90°). Additionally, a 124-slice high-resolution T1-weighted anatomical image was collected using spoiled gradient-recalled acquisition in steady-state imaging (TR = 5 ms, TE = 1.8 ms, flip angle = 15, FOV = 25−26 cm, slice thickness = 1.2 mm).

### fMRI Preprocessing

We used the Optimization of Preprocessing Pipelines for NeuroImaging software (OPPNI, (Churchill et al., 2012; Churchill Nathan W. et al., 2011) which perform automated processing and quality assessment of the fMRI data, using a combination of freeware packages from AFNI (https://afni.nimh.nih.gov) and FSL (https://www.fmrib.ox.ac.uk/fsl) along with custom algorithms. Processing included rigid-body motion correction (AFNI *3dvolreg*), slice-timing correction (AFNI *3dTshift*), spatial smoothing with 6mm FWHM Gaussian kernel (AFNI *3dmerge*), along with motion parameter regression (regressing out motion PCs that account for >85% of head movement parameters variance) and temporal detrending (Legendre polynomials of order 0 to 4) on the resting-state time-series. The data-driven PHYCAA+ algorithm (Churchill & Strother, 2013) was used to estimate and remove physiological noise components. We used a linear filter to suppress BOLD frequencies above 0.10 Hz. Finally, spatial normalization to group template was performed as follows: FSL *flirt* was used to compute the rigid-body transform of the mean functional volume for each participant in a session to their T1-weighted anatomical scan, along with the 12-parameter affine transformation of the T1 image for the participant to the MNI152 template. The transformation matrices were then concatenated, and the net transform applied to the fMRI data.

### Statistical Analysis

Contrast Partial Least Squares (PLS; (Krishnan, Williams, McIntosh, & Abdi, 2011; McIntosh & Lobaugh, 2004) analysis was used to identify the relationship between the set of behavioral variables organized as group-by-time blocks (results section 1), as well as the functional connectivity of the segmented brain regions from the whole brain with the group-by-time blocks (results section 2). To reduce the number of possible connections in the entire voxel space, i. e., C(102400, 2) = 5*10^9^, the whole brain for each subject was first segmented into 116 regions based on the Automated Anatomical Labeling (AAL) atlas (Tzourio-Mazoyer et al., 2002). This yields C(116, 2) = 6670 unique pairwise connections between AAL regions shown in Figure 1A as the sub-diagonal elements of the 116*116 correlation matrix for each subject at each time. The PLS technique allows for identification of significant relationships between two sets of variables (for example functional connectivity values and treatment levels in the study, e. g. see (Berman, Misic, et al., 2014; Cloutier, Li, Mišić, Correll, & Berman, 2017)) in a data-driven manner. The PLS implementation software was downloaded from Randy McIntosh‘s lab at: https://www.rotman-baycrest.on.ca/index.php?section=84. The version of PLS that we implemented was created on 05-JAN-2005 by Jimmy Shen and updated as part of Pls.zip: 16-MAY-2012.

In PLS, the goal of the analysis is to find weighted patterns of the original variables in the two sets (termed “latent variables” or “LVs”) that maximally co-vary with one another. In contrast PLS, these LVs represent a differentiation between levels of experimental design (i. e., three timepoints: pre-treatment, during treatment and post-treatment and three groups: chemotherapy, non-chemotherapy breast cancer, and aged-matched healthy controls in this study) interpreted as a contrast with 3*3 = 9 levels, as well as a spatial pattern of voxel activity that supports that contrast (in the case when the other set of variables is BOLD timeseries) or a weighted sum of the behavioral variables (when the other set of variables is behavioral variables). In the current study there are a total of 186 brain measurements, as well as 186 measurements for every behavioral variable (3 times for each participant in each group). These 186 measures are averaged across groups and time to create the **X** matrix, yielding an **X** of size 9 * 6670 and 9*11 for brain connections and behavioral variables, respectively. The **Y** matrix is the 9*9 experimental design matrix contrasting groups and times. PLS is computed via singular value decomposition (SVD; (Eckart & Young, 1936). The covariance between the two data sets **X** (e.g. mean-centered brain matrix of size 9 × 6670 connections or z-scored behavioral matrix of size 9 × 11 behavioral variables) and **Y** (e.g. experimental design matrix of size 9 × 9 group-by-time contrasts) is computed (**X′Y**) and is subjected to the SVD:

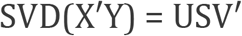

Resulting in a set of orthonormal matrices **U** and **V** (termed left and right singular vectors, respectively), as well as a diagonal matrix **S** of singular values. The number of LVs from the analysis is equal to the smallest rank of its constituent matrices (the rank of the covariance matrix **X′Y**, which is equal to 9, the degrees of freedom in the experimental design in the current study). The ith LV is comprised of a triplet of ith left singular vector, the ith right singular vector, and the ith singular value. The left and right singular vectors provide weights (or “saliences”) for the two sets (behavioral variables and treatment levels or voxels and treatment levels), respectively. The scalar singular value is proportional to the “crossblock covariance” between **X** and **Y** captured by the LV, and is naturally interpreted as the effect size of this statistical association.

The LVs are linear combinations of behavioral variables or voxel activities across the whole brain whose combinations are differentially instantiated for different groups * time cells. The left singular vector in the current study contains the contrast loadings for groups * time and the right singular vector contains the loadings for behavioral measures (Results section 1) or saliences of connectivity between brain regions (Results section 2). Multivariate methods applied to fMRI data offer a novel opportunity to discover meaningful associations between distributed patterns of brain activities and experimental conditions in a single statistical model, as opposed to univariate methods where conditions are regressed on every voxel or cluster of voxels separately. To test the significance of each LV, a set of 1000 covariance matrices were generated by randomly permuting condition labels for the **X** variables (brain or behavioral measures set). These covariance matrices embody the null hypothesis that there is no relationship between **X** and **Y** variables. They were subjected to SVD as before resulting in a null distribution of singular values. The significance of the original LV was assessed with respect to this null distribution. The *P* value was estimated as the proportion of the permuted singular values that exceed the original singular value.

The reliability with which each functional connection or behavioral variable contributes to the overall multivariate pattern was determined with bootstrapping. A set of 1000 bootstrap samples were created by re-sampling subjects with replacement within each group*time cell (i.e. preserving condition labels). Each new covariance matrix was subjected to SVD as before, and the singular vector weights from the resampled data were used to build a sampling distribution of the saliences from the original data set. The purpose of a constructed bootstrapped sampling distribution is to determine the reliability of each salience (i.e. saliences that are highly dependent on which participants are included in the analysis will have wide distributions).

For the brain connections, a single index of reliability (termed “bootstrap” ratio, or “Z_BR_”) was calculated by taking the ratio of the salience to its bootstrap estimated standard error. A Z_BR_ for a given connection is large when the connection has a large salience (i.e. makes a strong contribution to the LV) and when the bootstrap estimated standard error is small (i.e. the salience is stable across many resamplings). Here, connections with Z_BR_ > 3 or Z_BR_ < −3 were selected as showing reliable increase or decrease in functional connectivity, respectively (equivalent to p~0.0025, 2-tailed, under normal distribution assumptions). The process of applying PLS to relate the resting-state functional connectivity matrices to the groups*time contrast is described in Figure 1.

The process of applying PLS to the behavioral variables with the group*time contrast is similar, except that the matrix fed into the SVD contains the behavioral values instead of correlations between BOLD response of different brain regions. This would mean that each row vector for a participant at a timepoint is their 1*11 behavioral vector instead of 1*6670 connectivity vector. These row vectors are then stacked to create the behavioral matrix that will be used in the SVD (equivalent to the stacked connectivity matrix shown in Figure 1 section B).

**Figure 1.**
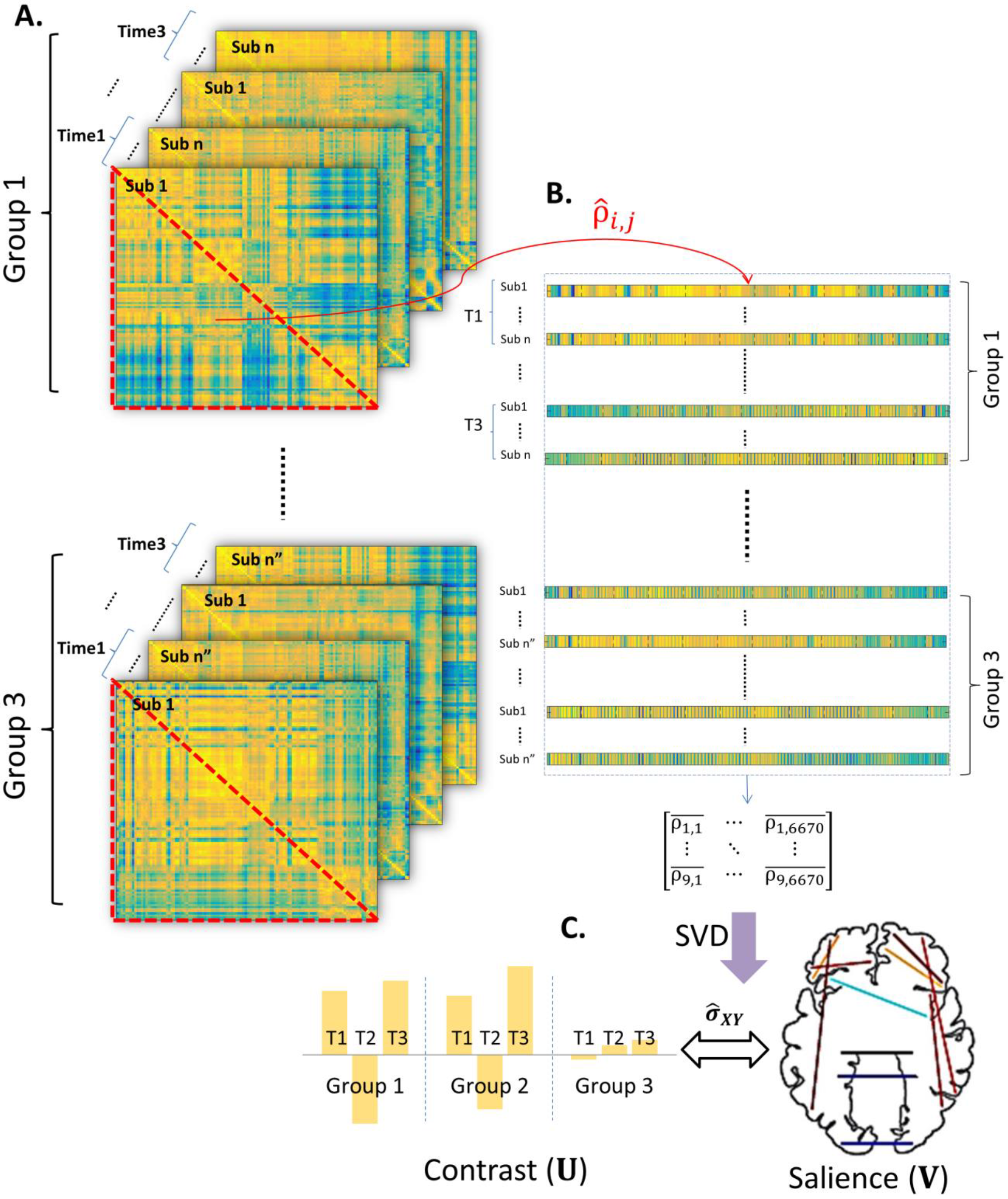
Schematic of contrast PLS for the brain data. **A.** Functional connectivity matrices: To reduce the number of possible connections in the all voxels space, i. e., C(102400, 2) = 5*10^9^, the whole brain for each subject was first segmented into 116 regions based on the Automated Anatomical Labeling (AAL) atlas (Tzourio-Mazoyer et al., 2002). This yieldsC(116, 2) = 6670 connections between AAL regions shown as a symmetric 116*116 correlation matrix for each subject at each time. **B.** The elements below main diagonal of connectivity matrices (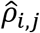 shows the Pearson correlation between BOLD time-series of two regions) are reshaped into a one-dimensional vector of size 1*6670 and stacked with the other participants within the same time, and then nested within the group. Then, the resulting matrix that contains all functional connectomes of all subjects averaged across times and groups and procedures described in the methods involving Singular Value Decomposition is applied to it. **C.** The resulting left singular vector (**U**, experimental conditions contrast) and right singular vector (**V**, brain connectivity salience) are shown, with the proportion of total covariance the LV is accounting for (Cross-block 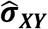) calculated as its squared singular value divided by sum of squares of singular values in the **S** matrix. The confidence intervals for loadings and bootstrap ratios for saliences are calculated using non-parametric (re-sampling) procedures described in the methods.

### Mediation Analysis

The mediations were implemented using R package ‘mediation’ (Tingley, Yamamoto, Hirose, Keele, & Imai, 2014) with quasi-Bayesian confidence intervals. The functional connectivity pattern associated with treatment and recovery was first aggregated for each participant at each scanning session. Because all the saliences are positive in the LV, this aggregation was done simply by summing up the Fisher’s z-transformed correlations between the regions that showed reliable connectivity change (i. e., the parietal-frontal connections with Z_BR_ > 3 from Figure 3). Alternatively, brain data from each subject can be projected on the singular vectors to get one aggregated score for the 214 saliences for each subject. The two methods yielded very similar results with aggregated measures from the two methods being correlated with r = .944. Hence, we are reporting the aggregated measure based on the first method described above as it is more intuitive. For simplicity, this will be referred to as the Aggregated Parieto-Frontal (APF) connectivity.

## Results

### Behavioral Results

We examined the primary time by group (M0, M1, and M7 by CT, nonCT, and HC) contrast, i.e., the one that covaried maximally with the behavioral measures, using the Partial Least Squares (PLS) multivariate analysis procedure (see Methods: *Statistical Analysis*). This latent variable is shown in Figure 2, the subsequent LVs did not pass the permutation null threshold of p < 0.01 and will not be discussed further. The Cross-block covariance between this emerged experimental condition contrast (Figure 2, top) and cognitive health measures (Figure 2, bottom) is relatively large (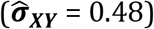), showing that the behavioral variables we measured as indicators of cognitive health can notably signify the status change of the patients pre-to post-treatment and recuperation.

The results show that the CT group at M1 is strongly affected by their treatment as exhibited in that group having more fatigue, depression, and physical burdens (BCPT), as well as lower self-reported attention (AFI) and lower performance on digit spans (DSB and DSF) compared to their baseline state at M0 (indicated by red line in Figure 2 top panel), and recovery state at M7 (indicated by green line in Figure 2 top panel). This decrease in objective cognitive performance and subjective emotional and cognitive well-being (henceforth referred to as decrease in cognitive health) at M1 compared to M0 and M7 is much smaller for the non-CT group and not present for the HC group. Therefore, we will be referring to this latent variable as chemotherapy treatment’s effect (detrimental effect M0 to M1, red) plus recuperation’s effect (beneficial effect M1 to M7, green) on cognitive health.

**Figure 2.**
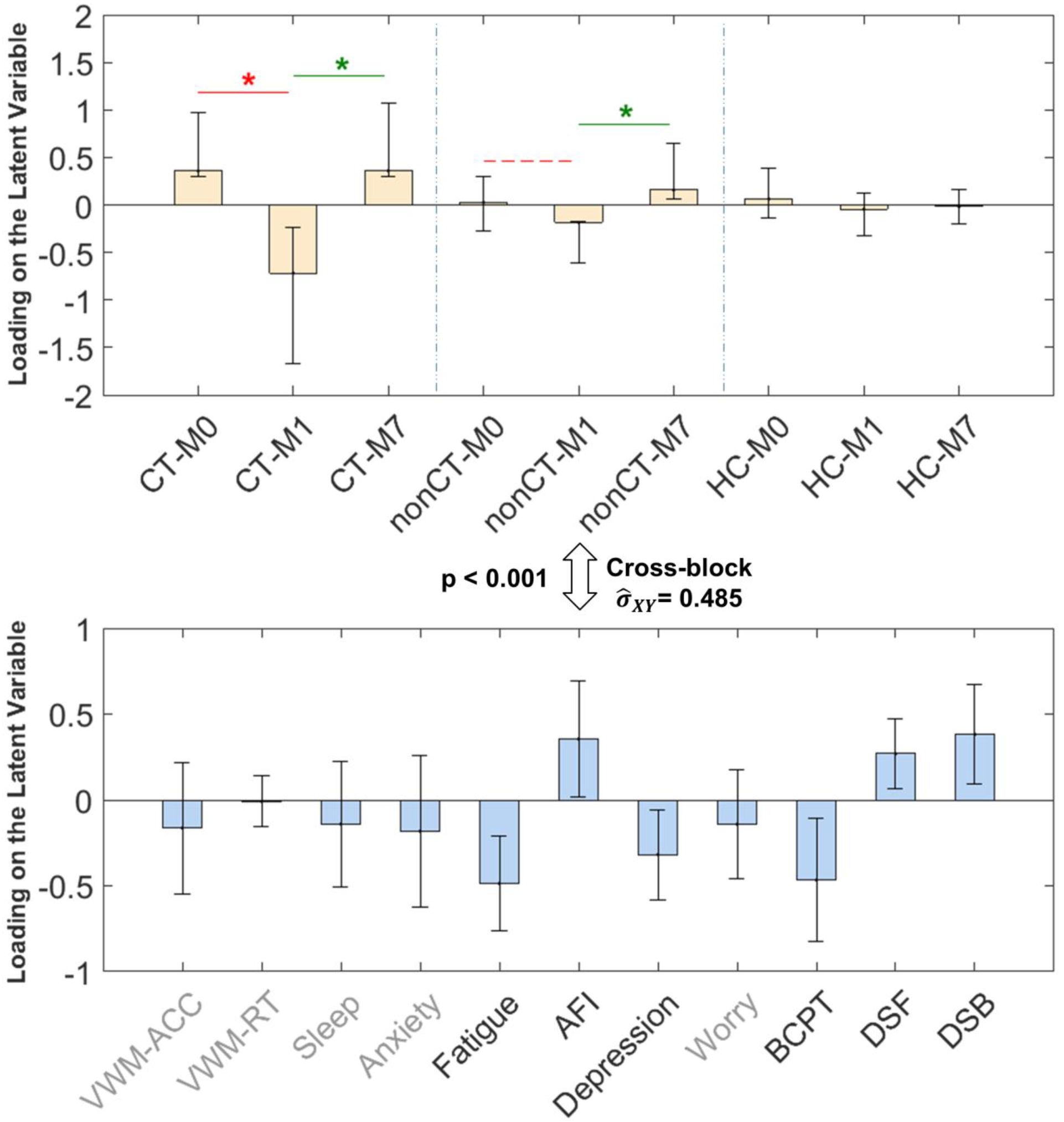
PLS results (LV1) relating experimental conditions (group * time, top panel) to the behavioral measures (bottom panel). Errorbars show 95% confidence intervals as indicated by bootstrapping. Cross-block covariance 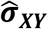 shows the cross-covariance between the two sets of data. The magnitude of loading for each variable shows its contribution to the LV. The red line with * indicates statistically significant (p < .001) treatment effect between M1 and M0 for the CT group. The green lines with * indicate significant (p < 0.001) recuperation effects between M1 and M7 for CT and non-CT groups.Grayed-out variable names show non-significant loading (p > 0.05) for the behavioral variable in the LV.

### Resting-state functional connectivity results

Next, PLS was used to find the optimum time by group contrast that maximally covaried with changes in functional connectivity between brain regions (see Methods). Here, again, the first latent variable is the only one that passed the permutation null threshold of p < 0.01 and therefore the subsequent LVs will not be discussed further.

The results, shown in Figure 3, indicate a relatively distributed pattern of functional connectivity mostly in parietal and frontal regions of the brain that decreases in connectivity strength at M1 compared to M0 and then increases again at M7 for patient groups (especially the CT group). There are 214 reliable connections in the pattern (Z_BR_ > 3) which includes ~3% of all the possible connections (See *Supplementary Results* for the list of the connections). Importantly, the emerged group-by-time contrast resembles the one previously found for behavioral measures (although not identical), hence we will refer to this latent variable as chemo treatment plus recuperation’s effects on functional connectivity. To emphasize the similarity between this LV and the one regarding cognitive health from the previous analysis, we again indicated the M0 to M1 contrast by red and the M1 to M7 contrast by green lines in Figure 3, respectively.

The proportion of covariance accounted for by this LV is also similar to that of the cognitive health measures (Cross-block 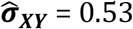 compared to .48 from the behavioral LV) which shows that this resting-state functional connectivity pattern could potentially be a neural signal that is as consistent as the cognitive health measures in capturing the effects of cancer treatment (especially chemotherapy) and recovery on patients.

**Figure 3.**
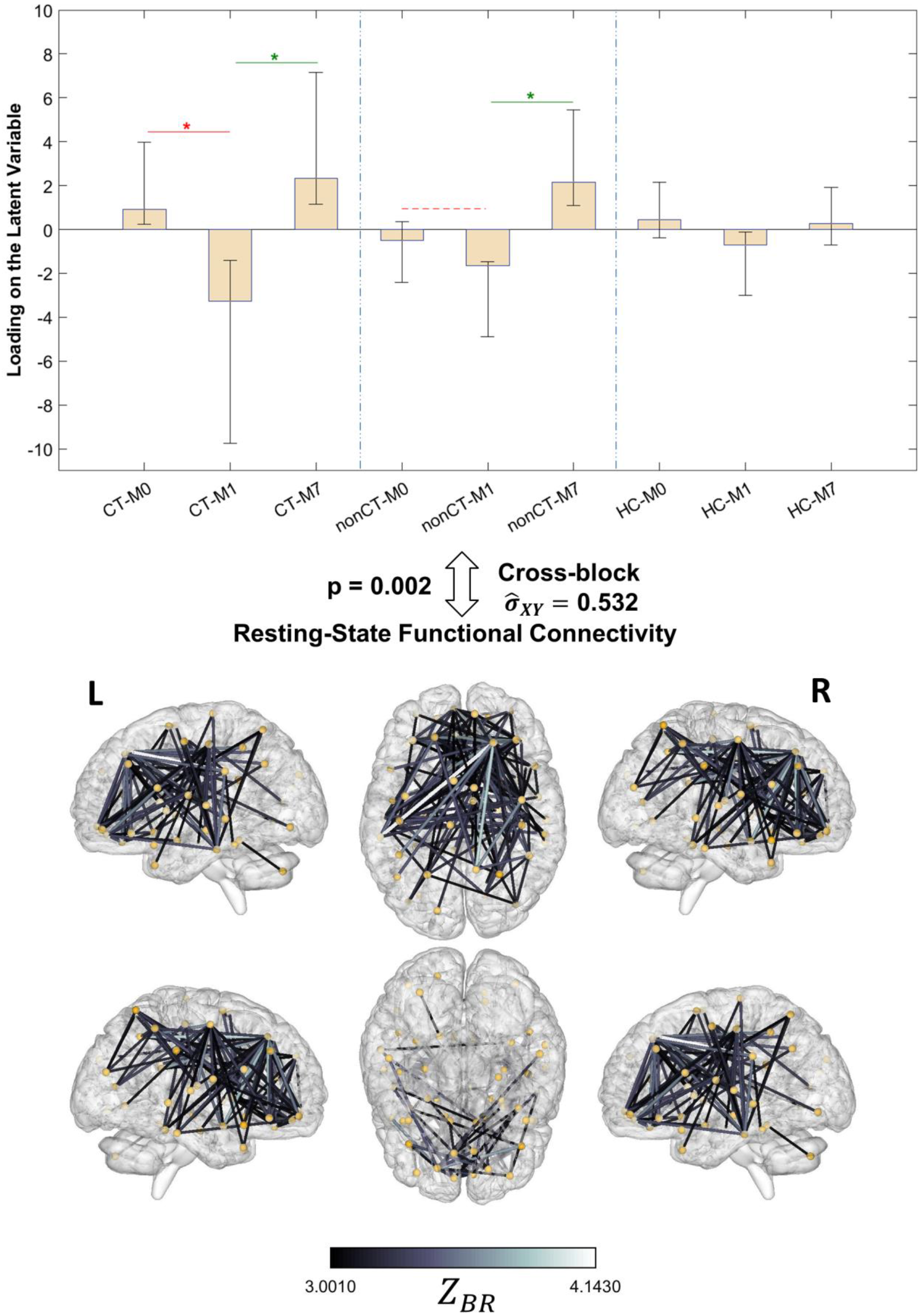
PLS results (LV1) relating experimental conditions (group * time, top panel) to brain functional connectivity (bottom panel). Errorbars show 95% confidence intervals as indicated by bootstrapping. Cross-block 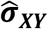 shows the proportion of covariance between the two sets explained by this LV. All edges in bottom panel show connections with Bootstrap ratio Z_BR_ values above 3 indicating reliable increase in connectivity strength, with lighter colors having greater _BR_ values. There were no connections with Z_BR_ < −3, indicating lack of any connections that reliably decreased for this latent variable. The red line with * in the top panel indicates statistically significant (p < .001) treatment effect between M1 and M0 for the CT group. The green lines with * in the top panel indicate significant (p < 0.001) recuperation effects between M1 and M7 for CT and non-CT groups.

### Relationship between cognitive health and functional connectivity measures

The similarity between the experimental conditions (time-by-group) LVs resulting from the two previous analyses (top panels in Figures 2 and 3, r = 0.903, Pearson correlation) suggests a potential relationship between the parieto-frontal functional connectivity pattern found in the brain connectivity LV (Figure 3 bottom panel) and the dip in cognitive health evident in the behavioral LV (Figure 2 bottom panel, i. e., more fatigue, depression, BCPT, lower AFI, DSB and DSF). In other words, both LVs could reflect the same phenomenon but in different domains (BOLD vs. behavior). Statistical evidence for this “common cause” hypothesis requires that shared covariance between the brain connectivity and experimental conditions overlaps with the shared covariance between the behavioral measures and experimental conditions.

To test this possibility, we first calculated an aggregated parieto-frontal (APF) measure of the functional connectivity pattern for each subject (see methods: *Mediation Analysis*). Next, we tested two alternative causal mediation models (Figure 4) to assess A) whether the APF connectivity disruption occurs, at least partially, as a result of cognitive health deficits (i. e., the LV consisted of higher fatigue, depression, BCPT, and lower AFI, DSB, and DSF), or alternatively B) whether the cognitive health deficits are partially caused by the APF connectivity disruption.

**Figure 4.**
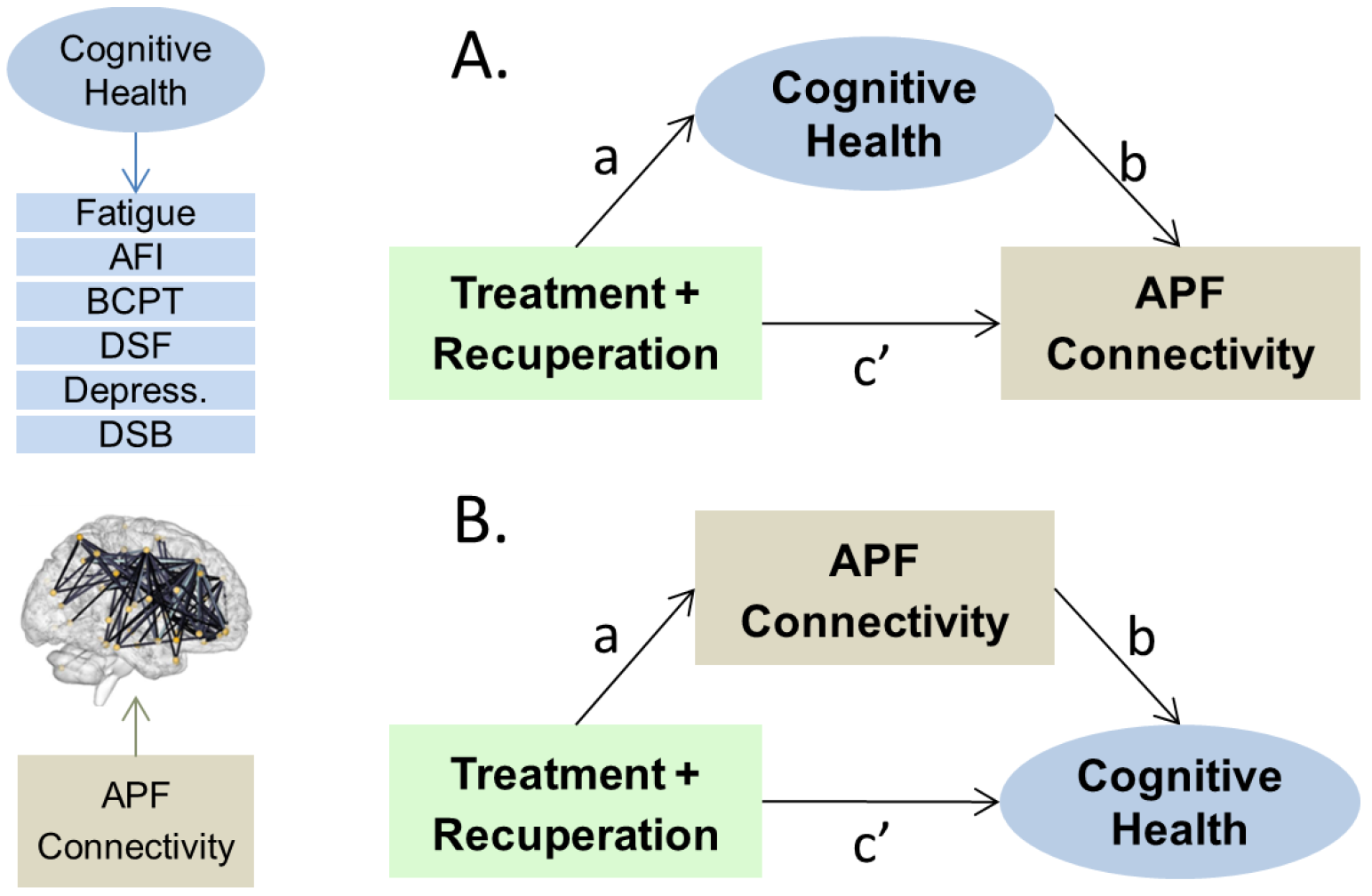
Two alternative causal mediation models. The panel on the left shows the constituent variables for cognitive health (top) and Aggregated Parieto-Frontal connectivity (bottom) latent variables, respectively. Treatment + Recuperation is simple contrast coding of M1 vs. M0 and M7for patient groups. **A.** Model used for testing if the chemotherapy + recuperation effects on functional connectivity in the parieto-frontal connectome is (partially) mediated by the cognitive health of the patients. **B.** Model used for testing if the chemotherapy + recuperation effects on cognitive health of the patients is (partially) mediated by functional connectivity in the parieto-frontal connectome. For both models the a*b path shows the mediated effect and the c’ path is the direct effect.

The results of the mediation analyses showed no evidence for either cognitive health mediating the chemotherapy treatment’s effect on APF connectivity (Model A: a*b = 0.01, Quasi Bayesian 95% CI = [−0.07, 0.08], p = .93) or the APF connectivity mediating the effect of chemotherapy on cognitive health (Model B: a*b = 0.02, Quasi Bayesian 95% CI = [-0.14, 0.16], p = 0.82). However, the direct effect of chemotherapy treatment + recuperation (c′) is significant in predicting both APF connectivity (Model A: c’ = 0.56, Quasi Bayesian 95% CI = [0.19, 0.91], p <0.01) and cognitive health (Model B: c’ = 0.79, Quasi Bayesian 95% CI = [0.18, 1.41], p <0.01) in both models. This implies that two different aspects of the chemotherapy process are responsible for the decrease in functional connectivity in the parietal-frontal connectome and increased distress and cognitive complaints after treatment.

## Discussion

In the current study, we first showed a deterioration in measures of cognitive and emotional well-being of breast cancer patients immediately after chemotherapy that they subsequently recovered from by 7 months after treatment. Specifically, the patients report more fatigue, depression, physical burdens (BCPT), and poorer attentional function (AFI), and perform worse on objective cognitive tasks (DSB and DSF) compared to their state before adjuvant therapy and 7 months after therapy. Parallel to this cognitive health deficit and recovery, we found an extensive, unidirectional change (decrease) in brain connectivity of the patients, where resting-state BOLD functional connectivity in parietal and frontal brain regions was disrupted as an apparent result of treatment (much stronger with chemotherapy but also present for non-chemotherapy treatments such as radiation therapy). Finding a single significant (high-variance) latent variable from the PLS analysis in the brain connectivity and behavior domains, as opposed to multiple non-primary LVs, indicates a robust response pattern across individuals in a single network of brain areas and set of behavioral variables, respectively.

Interestingly, similar to the cognitive health deficits, the functional connectivity disruption pattern also abates by 7 months. Despite the similarity in the longitudinal dynamics of the behavioral changes and brain connectivity disruption, the latter seems to emerge as a chemotherapy treatment symptom separate from the cognitive health deficits (i. e., neither caused by them nor causing them in a statistically significant manner as indicated by the results of our two mediation models). It is possible that these resting-state results reflect more basic physiological changes that may be challenging to measure with psychological tests. Our measurements of cognitive health cover a comprehensive spectrum of cognitive performance and emotional health. Admittedly, however, there could be other dimensions of cognitive functioning tied to prefrontal and posterior parietal cortices that the current study did not measure (such as complex decision making, reasoning, rumination, etc.) and are affected by the connectivity disruption reported here (or are responsible for the disruptions). Further research is required to pinpoint the mechanism through which chemotherapy is resulting in decreased resting-state BOLD connectivity. Until then, interventions aiming to improve the quality of life for cancer patients need to be directed towards improving behavioral performance and self-perceived well-being of the patient, especially during the first month after adjuvant chemotherapy.

The current study’s results imply that clinicians and medical professionals can utilize and apply resting-state functional connectivity in clinical populations more broadly and as a source of non-redundant information about patients, as it serves a complementary signal to the psychometric measures. While resting state functional connectivity has previously been used in clinical research and assessment of patient status (see for example (Fox & Greicius, 2010; Greicius, 2008), there are limited studies that directly compare its predictive power compared to that of more convenient ‘pencil and paper’ behavioral variables in clinical populations (Iordan et al., 2018; Kaiser, Andrews-Hanna, Wager, & Pizzagalli, 2015). We hope that these results help to motivate more inquiry about potential applications of resting-state functional connectivity signals to investigate side effects of intrusive, but life-saving, treatment procedures on brain function of clinical populations.

In conclusion, we have shown that resting-state functional connectivity in regions mostly within parieto-frontal network follows along the treatment and recovery dynamics of breast cancer treatment and recovery and explains as much variance in these treatment dynamics as a battery of behavioral self-report and objective measures of cognitive and physical well-being. We also demonstrated that this parieto-frontal resting state functional connectivity disruption fails to be proposed as a mechanistic explanation of the changes in cognitive well-being and the behavioral cognitive dysfunction of chemo patients. Interestingly and importantly, each measure explains unique variance in treatment dynamics. It will be important to understand what aspects of treatment and recovery are captured by the resting-state functional connectivity and what psychological factors are in fact related to it. More targeted support should be developed to understanding how to ameliorate the multifaceted side effects of cancer treatment and the ‘chemo-brain’.

## Acknowledgements

This work was supported in part by NIH grant (NIH-NINR-R01-NRO10939) to BC, a TKF foundation grant, Templeton Grant ID#: 37775 from the John Templeton Foundation (www.templeton.org) and a National Science Foundation (NSF) grant #1632445 to MGB.

## Supplementary Results

### Connections with Z_BR_ >3

**Figure S1.**
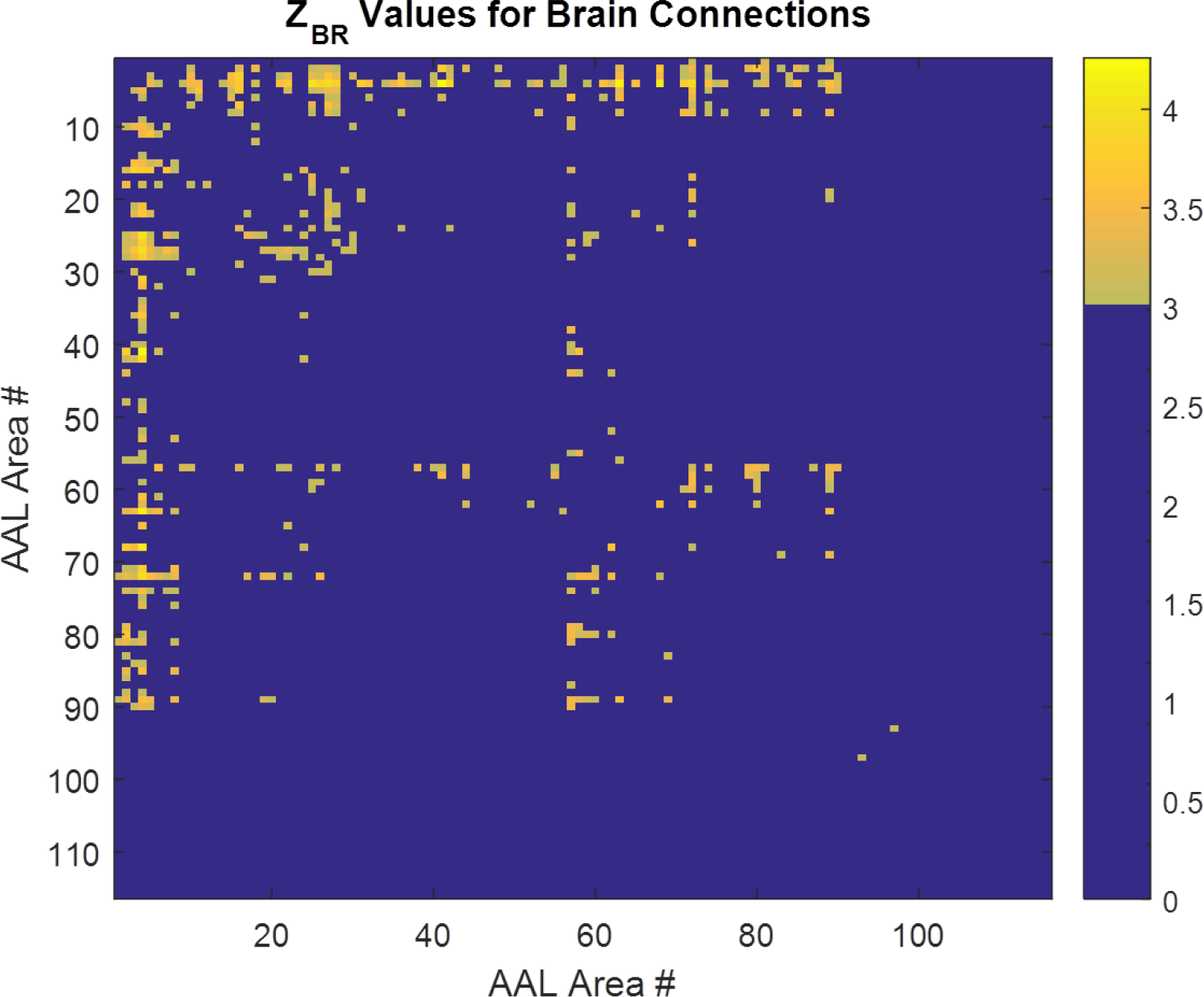
The connectivity matrix between AAL regions with bootstrap ratio Z values thresholded at 3. There are 214 connections that are listed below.

**Figure S2.**
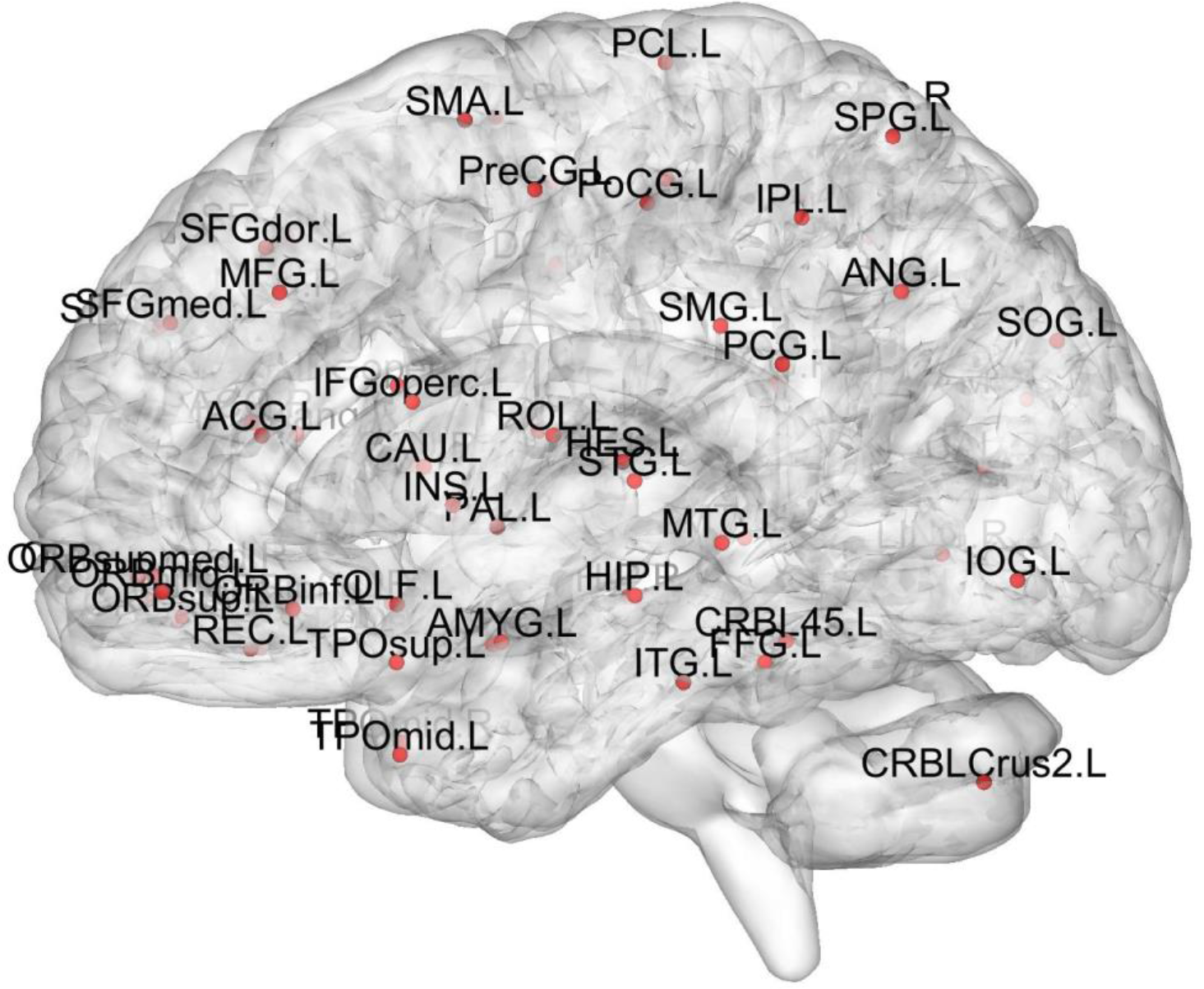
Guide to the labels of the AAL regions involved in the 214 connections with Z_BR_ > 3 that are listed below. Same labels as above with _R appended in the list below indicate corresponding contralateral regions in the right hemisphere.

### List of the connections

Precentral_L --------- Caudate_R50
Precentral_L --------- Temporal_Sup_L50
Precentral_L --------- Temporal_Inf_L50
Precentral_R --------- Frontal_Mid_Orb_R50
Precentral_R --------- Frontal_Inf_Orb_R50
Precentral_R --------- Rolandic_Oper_R50
Precentral_R --------- Frontal_Mid_Orb_L50
Precentral_R --------- Frontal_Mid_Orb_R50
Precentral_R --------- Rectus_L50
Precentral_R --------- Rectus_R50
Precentral_R --------- Amygdala_L50
Precentral_R --------- Amygdala_R50
Precentral_R --------- Calcarine_R50
Precentral_R --------- Lingual_R50
Precentral_R --------- Fusiform_R50
Precentral_R --------- SupraMarginal_L50
Precentral_R --------- Precuneus_R50
Precentral_R --------- Caudate_R50
Precentral_R --------- Putamen_R50
Precentral_R --------- Heschl_L50
Precentral_R --------- Heschl_R50
Precentral_R --------- Temporal_Sup_L50
Precentral_R --------- Temporal_Mid_L50
Precentral_R --------- Temporal_Mid_R50
Precentral_R --------- Temporal_Pole_Mid_R50
Precentral_R --------- Temporal_Inf_L50
Frontal_Sup_L --------- Frontal_Sup_Orb_L50
Frontal_Sup_L --------- Frontal_Mid_Orb_R50
Frontal_Sup_L --------- Frontal_Inf_Orb_L50
Frontal_Sup_L --------- Frontal_Inf_Orb_R50
Frontal_Sup_L --------- Olfactory_R50
Frontal_Sup_L --------- Frontal_Mid_Orb_L50
Frontal_Sup_L --------- Frontal_Mid_Orb_R50
Frontal_Sup_L --------- Rectus_L50
Frontal_Sup_L --------- Rectus_R50
Frontal_Sup_L --------- Insula_R50
Frontal_Sup_L --------- Cingulum_Post_R50
Frontal_Sup_L --------- ParaHippocampal_R50
Frontal_Sup_L --------- Amygdala_R50
Frontal_Sup_L --------- Fusiform_R50
Frontal_Sup_L --------- SupraMarginal_L50
Frontal_Sup_L --------- Precuneus_R50
Frontal_Sup_L --------- Caudate_L50
Frontal_Sup_L --------- Caudate_R50
Frontal_Sup_L --------- Temporal_Sup_L50
Frontal_Sup_L --------- Temporal_Pole_Sup_R50
Frontal_Sup_L --------- Temporal_Inf_R50
Frontal_Sup_R --------- Frontal_Sup_Orb_L50
Frontal_Sup_R --------- Frontal_Mid_Orb_L50
Frontal_Sup_R --------- Frontal_Mid_Orb_R50
Frontal_Sup_R --------- Frontal_Inf_Oper_L50
Frontal_Sup_R --------- Frontal_Inf_Tri_R50
Frontal_Sup_R --------- Frontal_Inf_Orb_L50
Frontal_Sup_R --------- Frontal_Inf_Orb_R50
Frontal_Sup_R --------- Rolandic_Oper_R50
Frontal_Sup_R --------- Olfactory_L50
Frontal_Sup_R --------- Olfactory_R50
Frontal_Sup_R --------- Frontal_Mid_Orb_L50
Frontal_Sup_R --------- Frontal_Mid_Orb_R50
Frontal_Sup_R --------- Rectus_L50
Frontal_Sup_R --------- Rectus_R50
Frontal_Sup_R --------- Cingulum_Ant_L50
Frontal_Sup_R --------- Cingulum_Ant_R50
Frontal_Sup_R --------- Cingulum_Mid_R50
Frontal_Sup_R --------- Cingulum_Post_L50
Frontal_Sup_R --------- Cingulum_Post_R50
Frontal_Sup_R --------- Hippocampus_L50
Frontal_Sup_R --------- Hippocampus_R50
Frontal_Sup_R --------- ParaHippocampal_R50
Frontal_Sup_R --------- Amygdala_L50
Frontal_Sup_R --------- Amygdala_R50
Frontal_Sup_R --------- Lingual_R50
Frontal_Sup_R --------- Occipital_Sup_L50
Frontal_Sup_R --------- Occipital_Inf_L50
Frontal_Sup_R --------- Fusiform_L50
Frontal_Sup_R --------- Parietal_Inf_L50
Frontal_Sup_R --------- Parietal_Inf_R50
Frontal_Sup_R --------- SupraMarginal_L50
Frontal_Sup_R --------- Angular_L50
Frontal_Sup_R --------- Precuneus_R50
Frontal_Sup_R --------- Caudate_L50
Frontal_Sup_R --------- Caudate_R50
Frontal_Sup_R --------- Putamen_R50
Frontal_Sup_R --------- Pallidum_L50
Frontal_Sup_R --------- Pallidum_R50
Frontal_Sup_R --------- Heschl_R50
Frontal_Sup_R --------- Temporal_Sup_L50
Frontal_Sup_R --------- Temporal_Pole_Sup_R50
Frontal_Sup_R --------- Temporal_Mid_L50
Frontal_Sup_R --------- Temporal_Pole_Mid_R50
Frontal_Sup_R --------- Temporal_Inf_L50
Frontal_Sup_R --------- Temporal_Inf_R50
Frontal_Sup_Orb_L --------- Frontal_Mid_Orb_R50
Frontal_Sup_Orb_L --------- Frontal_Inf_Oper_L50
Frontal_Sup_Orb_L --------- Frontal_Inf_Orb_L50
Frontal_Sup_Orb_L --------- Frontal_Inf_Orb_R50
Frontal_Sup_Orb_L --------- Olfactory_R50
Frontal_Sup_Orb_L --------- Frontal_Mid_Orb_L50
Frontal_Sup_Orb_L --------- Frontal_Mid_Orb_R50
Frontal_Sup_Orb_L --------- Rectus_L50
Frontal_Sup_Orb_L --------- Rectus_R50
Frontal_Sup_Orb_L --------- SupraMarginal_L50
Frontal_Sup_Orb_L --------- Caudate_R50
Frontal_Sup_Orb_L --------- Temporal_Inf_L50
Frontal_Sup_Orb_L --------- Temporal_Inf_R50
Frontal_Sup_Orb_R --------- Frontal_Inf_Oper_L50
Frontal_Sup_Orb_R --------- Frontal_Inf_Orb_L50
Frontal_Sup_Orb_R --------- Rolandic_Oper_R50
Frontal_Sup_Orb_R --------- Rectus_L50
Frontal_Sup_Orb_R --------- Rectus_R50
Frontal_Sup_Orb_R --------- Amygdala_L50
Frontal_Sup_Orb_R --------- Postcentral_L50
Frontal_Sup_Orb_R --------- SupraMarginal_L50
Frontal_Sup_Orb_R --------- Caudate_R50
Frontal_Mid_L --------- Frontal_Mid_Orb_R50
Frontal_Mid_L --------- Frontal_Inf_Orb_R50
Frontal_Mid_L --------- Frontal_Mid_Orb_L50
Frontal_Mid_L --------- Rectus_L50
Frontal_Mid_L --------- Rectus_R50
Frontal_Mid_L --------- Caudate_R50
Frontal_Mid_R --------- Frontal_Inf_Orb_R50
Frontal_Mid_R --------- Frontal_Mid_Orb_L50
Frontal_Mid_R --------- Rectus_L50
Frontal_Mid_R --------- Rectus_R50
Frontal_Mid_R --------- Cingulum_Post_R50
Frontal_Mid_R --------- Occipital_Inf_L50
Frontal_Mid_R --------- SupraMarginal_L50
Frontal_Mid_R --------- Caudate_L50
Frontal_Mid_R --------- Caudate_R50
Frontal_Mid_R --------- Putamen_R50
Frontal_Mid_R --------- Temporal_Sup_L50
Frontal_Mid_R --------- Temporal_Mid_L50
Frontal_Mid_R --------- Temporal_Inf_L50
Frontal_Mid_Orb_L --------- Postcentral_L50
Frontal_Mid_Orb_R --------- Rolandic_Oper_R50
Frontal_Mid_Orb_R --------- Insula_R50
Frontal_Mid_Orb_R --------- Postcentral_L50
Frontal_Inf_Oper_R --------- Rolandic_Oper_R50
Frontal_Inf_Orb_R --------- Frontal_Sup_Medial_R50
Frontal_Inf_Orb_R --------- Insula_L50
Frontal_Inf_Orb_R --------- Postcentral_L50
Rolandic_Oper_L --------- Olfactory_R50
Rolandic_Oper_L --------- Frontal_Mid_Orb_L50
Rolandic_Oper_L --------- Caudate_R50
Rolandic_Oper_R --------- Frontal_Mid_Orb_L50
Supp_Motor_Area_L --------- Rectus_L50
Supp_Motor_Area_L --------- Cingulum_Ant_L50
Supp_Motor_Area_L --------- Caudate_R50
Supp_Motor_Area_L --------- Temporal_Inf_L50
Supp_Motor_Area_R --------- Rectus_L50
Supp_Motor_Area_R --------- Cingulum_Ant_L50
Supp_Motor_Area_R --------- Caudate_R50
Supp_Motor_Area_R --------- Temporal_Inf_L50
Olfactory_L --------- Rectus_L50
Olfactory_L --------- Rectus_R50
Olfactory_L --------- Postcentral_L50
Olfactory_R --------- Frontal_Sup_Medial_R50
Olfactory_R --------- Rectus_L50
Olfactory_R --------- Rectus_R50
Olfactory_R --------- Postcentral_L50
Olfactory_R --------- Angular_L50
Frontal_Sup_Medial_L --------- Rectus_L50
Frontal_Sup_Medial_R --------- Frontal_Mid_Orb_L50
Frontal_Sup_Medial_R --------- Rectus_L50
Frontal_Sup_Medial_R --------- Rectus_R50
Frontal_Sup_Medial_R --------- Cingulum_Post_R50
Frontal_Sup_Medial_R --------- Amygdala_R50
Frontal_Sup_Medial_R --------- Precuneus_R50
Frontal_Mid_Orb_L --------- Insula_R50
Frontal_Mid_Orb_L --------- Parietal_Sup_L50
Frontal_Mid_Orb_L --------- Parietal_Sup_R50
Frontal_Mid_Orb_R --------- Rectus_R50
Frontal_Mid_Orb_R --------- Insula_R50
Frontal_Mid_Orb_R --------- Postcentral_L50
Frontal_Mid_Orb_R --------- Parietal_Sup_L50
Frontal_Mid_Orb_R --------- Caudate_R50
Rectus_L --------- Insula_L50
Rectus_L --------- Insula_R50
Hippocampus_R --------- Postcentral_L50
ParaHippocampal_R --------- Postcentral_L50
Amygdala_L --------- Postcentral_L50
Amygdala_L --------- Postcentral_R50
Calcarine_R --------- Postcentral_L50
Calcarine_R --------- Postcentral_R50
Calcarine_R --------- Parietal_Inf_R50
Occipital_Mid_R --------- Parietal_Inf_R50
Fusiform_L --------- Postcentral_R50
Fusiform_R --------- SupraMarginal_L50
Postcentral_L --------- Caudate_R50
Postcentral_L --------- Putamen_R50
Postcentral_L --------- Heschl_L50
Postcentral_L --------- Heschl_R50
Postcentral_L --------- Temporal_Sup_L50
Postcentral_L --------- Temporal_Pole_Mid_L50
Postcentral_L --------- Temporal_Inf_L50
Postcentral_L --------- Temporal_Inf_R50
Postcentral_R --------- Caudate_R50
Postcentral_R --------- Heschl_L50
Postcentral_R --------- Heschl_R50
Postcentral_R --------- Temporal_Inf_L50
Parietal_Sup_L --------- Caudate_R50
Parietal_Sup_L --------- Heschl_R50
Parietal_Sup_L --------- Temporal_Inf_L50
Parietal_Sup_R --------- Caudate_L50
Parietal_Sup_R --------- Caudate_R50
Parietal_Sup_R --------- Heschl_R50
Parietal_Sup_R --------- Temporal_Inf_L50
Parietal_Inf_R --------- Precuneus_R50
Parietal_Inf_R --------- Caudate_R50
Parietal_Inf_R --------- Heschl_R50
SupraMarginal_L --------- Temporal_Inf_L50
Paracentral_Lobule_L --------- Temporal_Pole_Sup_L50
Paracentral_Lobule_L --------- Temporal_Inf_L50
Cerebelum_Crus2_L --------- Cerebelum_4_5_L50

